# Steering Molecular Dynamics Simulations of Membrane-Associated Proteins with Neutron Reflection Results

**DOI:** 10.1101/823740

**Authors:** Bradley W. Treece, Frank Heinrich, Arvind Ramanathan, Mathias Lösche

**Affiliations:** Department of Physics, Carnegie Mellon University, Pittsburgh, PA 15213, USA; Department of Biomedical Engineering, Carnegie Mellon University, Pittsburgh, PA 15213, USA; The NIST Center for Neutron Research, National Institute of Standards and Technology, Gaithersburg, MD 20899, USA; Data Science and Learning, Argonne National Laboratory, Lemont, IL 60439, USA

**Keywords:** neutron reflection, protein structure, integrative modeling, steered molecular dynamics

## Abstract

We present a method to incorporate structural results from neutron reflectometry, a technique that determines interfacial structures such as protein-membrane complexes at a solid surface, into molecular dynamics simulations. By analyzing component volume occupancy profiles, which describe the one-dimensional distribution of a particular molecular component within an interfacial architecture, we construct a real-space constraint in the form of a biasing potential for the simulation that vanishes when the simulated and experimental profiles agree. This approach improves the correspondence between simulation and experiment, as shown for an earlier investigation where an NR-derived structure was well captured by an independent MD simulation, and may lead to faster equilibration of ensemble structures. We further show that time averaging of the observable when biasing with this approach permits fluctuations about the average, which are necessary for conformational exploration of the protein. The method described here also provides insights into systems that are characterized by NR and MD when the two show slight differences in their profiles. This is particularly valuable for studies of proteins at interfaces that contain disordered regions since the conformation of such regions is difficult to judge from the analysis of one-dimensional experimental profiles and take prohibitively long to equilibrate in simulations.

## 1. Introduction

Constrained computer simulations have been broadly used to identify models of molecular systems that are consistent with experimental observations.^1-3^ In molecular dynamics (MD) simulations, biasing potentials usually achieve this by tilting the energy landscape such that molecular conformations which comply with known experimental outcomes are favored in the development of molecular trajectories. A variety of approaches are routinely used to align molecular models in MD with X-ray or NMR structures by adding a bias to the intermolecular forces that drive the MD computation engine.^4^ However, such interferences can corrupt the statistical properties of fluctuations in the computational model in undesirable ways.^5^ This problem can be mitigated by constructing bias potentials from time-averaged ensembles of experimental observable(s) produced in the course of the simulation, as implemented in early implementations for the steering of MD simulations to reflect NMR cross-relaxation^6^ and paramagnetic relaxation results.^7^ Unperturbed distributions within an ensemble can be identified by minimally invasive biasing in accordance with the maximum entropy method,^8^ and this approach is equivalent to restrained-ensemble simulations, as shown by Roux and Weare.^5^ Biased simulations are thus expected to speed up equilibration of the system. To estimate the gains in efficiency of sampling equilibrium properties in biased simulations, one may evaluate time series of relevant observables to determine the time points in trajectories with the maximum number of uncorrelated samples in the series.^9^ Here, we develop a method for biasing MD simulations of protein complexes in lipid membranes toward experimental results from reflectometry measurements of substrate-supported model systems.^10^

Lipid bilayers are important intracellular structures that organize vital cellular activities in which membrane-associated proteins play critical roles. These can be firmly bound as transmembrane proteins or transiently interact with the bilayer interface as peripheral proteins. In either case, the membrane-bound structure of the protein can be quite distinct from its conformation in solution. Specular neutron reflection (NR) is the technique of choice for characterizing the structure of such proteins in the context of a lipid bilayer with molecular-scale resolution. We recently showed that planar lipid membranes, assembled on a substrate, provide an in-plane fluid matrix^11-12^ to which membrane proteins can be incorporated^13^ or peripherally bound,^14-17^ and characterized by NR with high resolution.^10,18^ This technology is unique among structural biology methods in that it is performed at room temperature under physiological conditions without any fixation because the bilayer structure, while in-plane fluid, is extraordinarily stable.^19^ Their resilience in this sample format, which we refer to as sparsely-tethered bilayer lipid membranes (stBLMs),^11^ permits sequential characterization of as-prepared lipid membranes under distinct solvent conditions such as different isotopic buffers, which is key to assessing their structure at high resolution.^10,20-22^ Subsequent investigation of the same sample after protein incorporation then enables one to determine the structure of a protein-membrane complex. ^13-15,23^ As the scattering does not depend on a periodic sample arrangement, (partially) disordered proteins may be characterized,^16^ and because proteins are associated with a single bilayer membrane that exposes an open face to a semi-infinite buffer reservoir, structural responses to extrinsic triggers can be recorded.^15^ Scattering from the protein-membrane structure in reflection mode is intrinsically selective to the interfacial structure,^24^ *i.e.* insensitive to the scattering from a potentially large concentration of protein that may remain dissolved in the adjacent buffer. This interfacial sensitivity permits the structural characterization of minute amounts of membrane-bound protein.^13^

The momentum transfer in a specular scattering experiment is strictly perpendicular to the bilayer and provides information about the interfacial structure in the form of one-dimensional scattering length density (SLD) profiles. The canonical resolution of such a profile in a single NR measurement is only on the order of 1 nm.^20^ However, contrast variation of the buffer and the measurement of a specific sample before and after protein association increases the information content dramatically.^25^ In addition, modern modeling techniques that combine atomistic protein structures with parameterized bilayer structures^26^ permit the localization of a protein in the membrane with a resolution of 1 Å.^13^ In combination, this methodology provides in-plane averaged material density distributions along the interfacial normal direction *z* – which we call component volume occupancy (CVO) profiles, *ρ*_cvo_(*z*) – that locate the protein within the bilayer with a precision that borders atomistic resolution.^10^ A more detailed account of interfacial scattering and its relation to MD simulations is provided as Supplemental Information.

Ultimately, molecular dynamics (MD) simulations of membrane-protein complexes are the method of choice to develop models of membrane-associated proteins on bilayers. In particular, MD can assist the interpretation of NR-derived structural information and access levels of detail inaccessible to NR. We showed previously that the joint interpretation of NR and MD data reveals minute structural adjustments of the PTEN tumor suppressor to the bilayer upon binding and identified lipid specific interactions of its binding domain.^18,27^ But there is an important caveat: obtaining meaningful correlations between MD-derived and NR-derived structures requires that the simulations sample equilibrium states. It has been extensively documented that equilibrium conditions in atomistic MD simulations of bilayer systems are notoriously difficult to achieve^28-32^ because sampling a full ensemble of configurations in such large systems requires prohibitively long simulation time. Restraining the phase space of simulations to conformational ensembles that reproduce experimental observables, generally referred to as *integrative modeling*,^33-34^ is a mature concept in electron microscopy,^33^ crystallography,^35-37^ NMR,^38-39^ and SAXS.^40-41^

Here, we configure and test a technique to steer MD simulations of proteins in or near bilayer membranes using *ρ*_cvo_(*z*) from NR measurements as a global restraint which biases toward conformational responses of the protein that match the experimental results. This is accomplished by a potential *U*_st_(*z*) proportional to the discrepancy between the CVO profiles derived from experiment (*ρ*_exp_(*z*)) and simulation (*ρ*_sim_(*z*)). It is adaptive, taking into account the dynamical nature of *ρ*_sim_(*z*), and vanishes upon agreement of the two profiles. Testing this method in a GROMACS implementation on two model systems, we fine-tune its strength relative to the interatomic potentials by steering a helical peptide to a predetermined position and orientation in solution. Two modes of determining *ρ*_sim_(*z*) are explored. It is observed that drawing a sample of *ρ*_sim_(*z*) instantaneously from the developing trajectory leads to a hard constraint that may not be desirable. However, if *ρ*_sim_(*z*) is determined as a running average of samples drawn along the developing trajectory, the position of the test structure is observed to be well constrained while fluctuation amplitudes are similar to those observed in a free simulation of the same system.

We then apply our insights from the peptide simulations to steered simulations of a well-studied, midsize protein near a membrane, for which NR data were previously interpreted with free MD simulations.^27^ We observe that all simulations, free and biased, lead to protein conformations on the membrane that are roughly consistent with the experimental results (as were the previous, more extensive free simulations). However, only simulation under moderate bias strength and continuously averaged feedback produced a trajectory in which all observables we analyzed converged efficiently within the 100 ns simulation time, according to the Chodera criterion.^9^ The applied bias strength was sufficiently gentle to not distort the folded domains of the protein; yet, simulations biased with continuously averaged feedback, apparently exploring broader ranges of the phase space, identified novel conformations for the long disordered C-terminal tail of the simulated protein that are plausible and may shed new light on the regulatory function of this protein segment.

## 2. Methods

The basic algorithm that biases the simulation toward an experimental result is bipartite, depending on whether *ρ*_exp_ and *ρ*_sim_ do or do not overlap. If profiles overlap, a steering potential is added to the molecular force field and drives a particular system component of interest – typically a protein – toward ensemble configurations for which 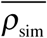, the averaged density, matches *ρ*_exp_. When profiles do not overlap, initially or in response to fluctuations mid-simulation, a global force on all its atoms attracts the steered component toward its desired position along *z* until profile overlap is reestablished.

### 2.1 Steering Potential

The profile *ρ*_cvo_ of a molecular configuration is the projection, on the direction perpendicular to the interface, *z*, of the in-plane averaged relative volume densities of suitably parsed molecular components (see Supplement). Such components can include ‘phospholipid chains’, ‘the protein’ or, in a distinct choice for parsing, ‘protein domain A’ and ‘protein domain B’. For the construction of a steering potential, only the relevant component is guided toward its experimentally determined position in the profile along *z*. For simplicity, assume that there is only one protein species in the experimental situation and that it is homogeneously labeled (hydrogenated or deuterated), so that its internal domains are not distinguished in an NR experiment. Its one-dimensional component density is

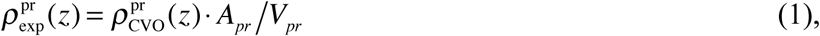

measured in nm^−1^, where *V*_*pr*_ is the volume of one molecule and *A*_*pr*_ is its projected area onto the interface.^*^ To suitably bias protein atoms, we determine representations of the corresponding volume density distribution from the simulation as

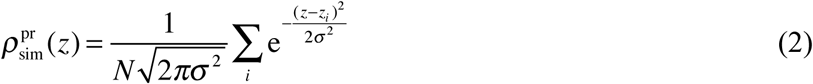

where *i* enumerates all *N* component atoms. This provides a steering potential,

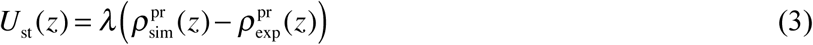

in which *λ* sets the energy scale (in kJ/mol or k_B_*T* per molecule). Because *ρ* is associated with an inverse length (Eq. (1)), we quote *λ* in k_B_*T*·nm. The supplemental potential *U* results in forces, *F*_*z*_ = – d*U*/d*z*, that attract protein atoms toward regions where 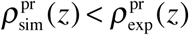, repel otherwise, and vanish at locations where the two densities match, as illustrated in Fig. 1 for the hypothetical case of a misaligned helical peptide in solution. In formal terms, Eq. (3) introduces a Lagrange multiplier that restricts the phase space to regions where the simulated density conforms to the one experimentally observed. In this formalism, deviations from the constraint are met with forces opposing them, and only when the system is in a region of phase space that is consistent with the constraint, does the term vanish from the equations of motion. However, in contrast to the Lagrangian analogy, *U*_st_(*z*) does not entirely prevent deviations from the constraint, as that would require infinite force.^5,42^

**Figure 1:**
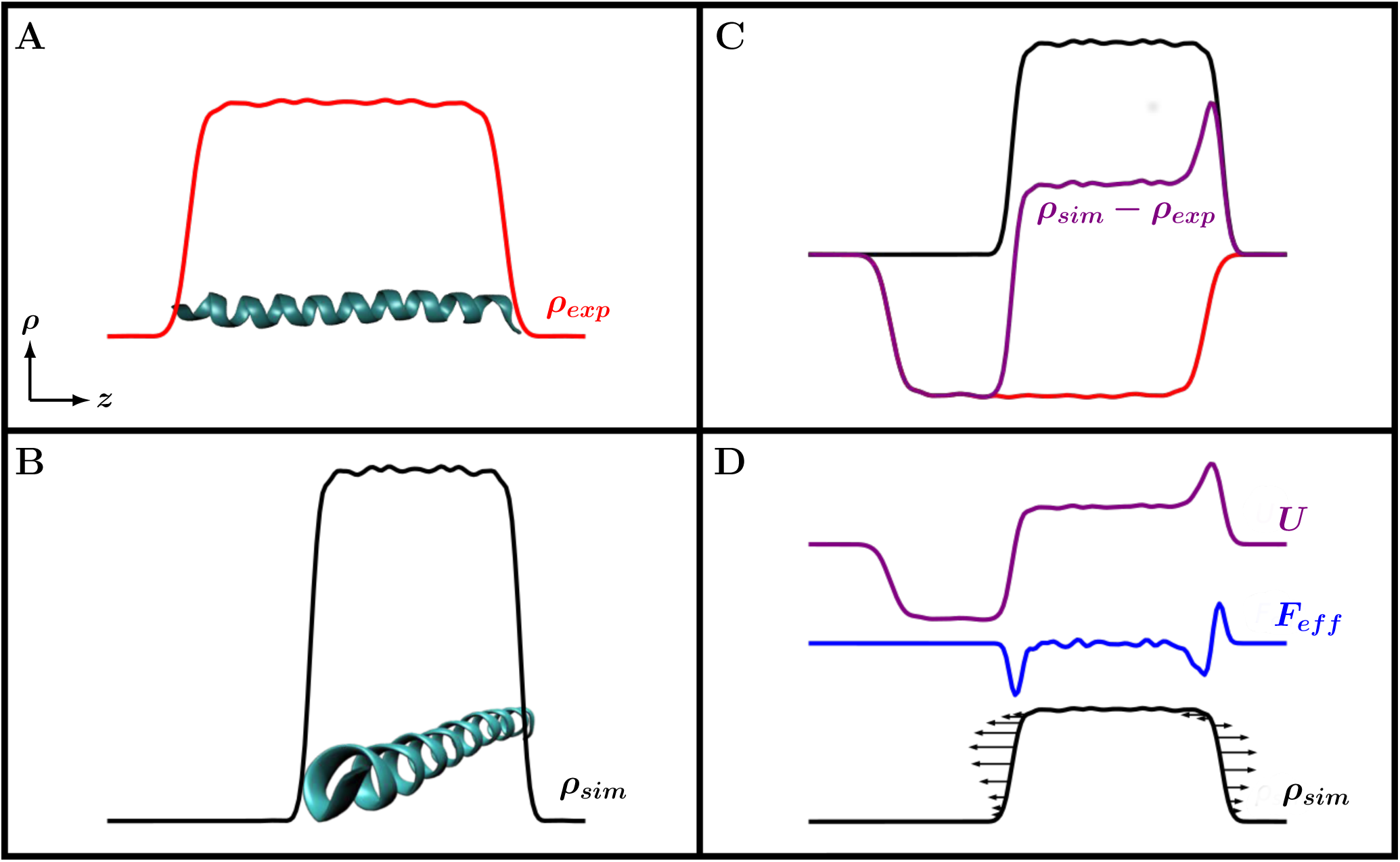
Input data structures and general strategy for the derivation of biasing potentials in MD simulations steered toward experimental NR results. (A) A fluctuating molecular structure gives rise to a one-dimensional profile *ρ*_exp_(*z*) such as those derived from NR (for details, see Fig. S1). In this exemplification, *ρ*_exp_(*z*) is derived from the unbiased simulation of an aligned peptide helix, described in section 3.1. (B) An instantaneous snapshot of the helix disoriented with respect to *z* gives rise to a profile *ρ*_sim_(*z*). (C) The steering potential *U* is proportional to the difference between *ρ*_sim_(*z*) and *ρ*_exp_(*z*). (D) The resulting forces and their impact on *ρ*_sim_(*z*) create a torque on the helix that enforces alignment with the target structure.

The feedback provided by 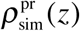 in Eq. (3) can be instantaneous: 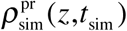 is determined at the time, *t*_sim_, when the subsequent step in the simulation in evaluated. Alternatively, it can be used as a time average,

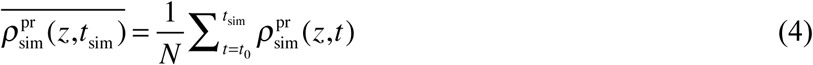

where *N* is the number of samples drawn from the trajectory between *t* = *t*_0_ and *t*_sim._ 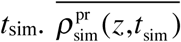 reflects the ensemble character of the sampled configuration in the course of the simulation and also may be better capable of capturing an actual experimental result in which membrane-bound proteins obtain locally distinct conformations over which an NR measurement averages in-plane.

To manage situations where 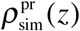 and 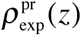 do not (sufficiently) overlap, we determine the two leading moments of their density functions, *µ*^(1)^ and *µ*^(2)^. If

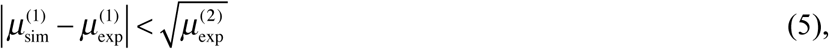

steering is provided according to Eq. (3). Conversely, if the profiles do not overlap, the simulation density is driven toward the experimental density by adding

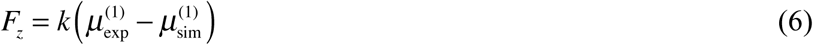

to the intrinsic forces in which *k* is scaled by *N*. In this case, the non-overlapping profiles are not included in the continuously accumulating average that determines 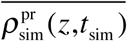.

To determine relative measures for how fast simulations loose their initial bias and approach equilibration under different conditions, we use the method of Chodera which selects that point in the time series of an observable that maximizes the number of uncorrelated samples in that series. This approach has been reported to yield robust estimates even if only a small number of independent trajectories are available for evaluation.^9^

### 2.2. Implementation and System Set-up

We implemented modifications to GROMACS 5.1.1 to compute and incorporate the potential according to Eq. (3) as presented in Fig. 2. The target distribution 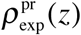 is determined from NR data and handed to the preprocessor together with a suitable value of *λ* and the initial configuration of the simulation to create the run file. At production runtime, forces are added to protein atoms by evaluating the gradients of the overall potential at their locations. These forces, along with molecular force field of the simulation, then determine the dynamics of the protein coordinates. The resulting coordinates are periodically retrieved to update 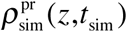 or 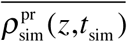. In the examples discussed below, this update occurs at 0.1 ps intervals with σ set to 1 Å and each atom’s distribution is truncated at 3σ for computational efficiency. Tuning of *λ* is explored in parallel simulation runs, and empirically optimized.

**Figure 2:**
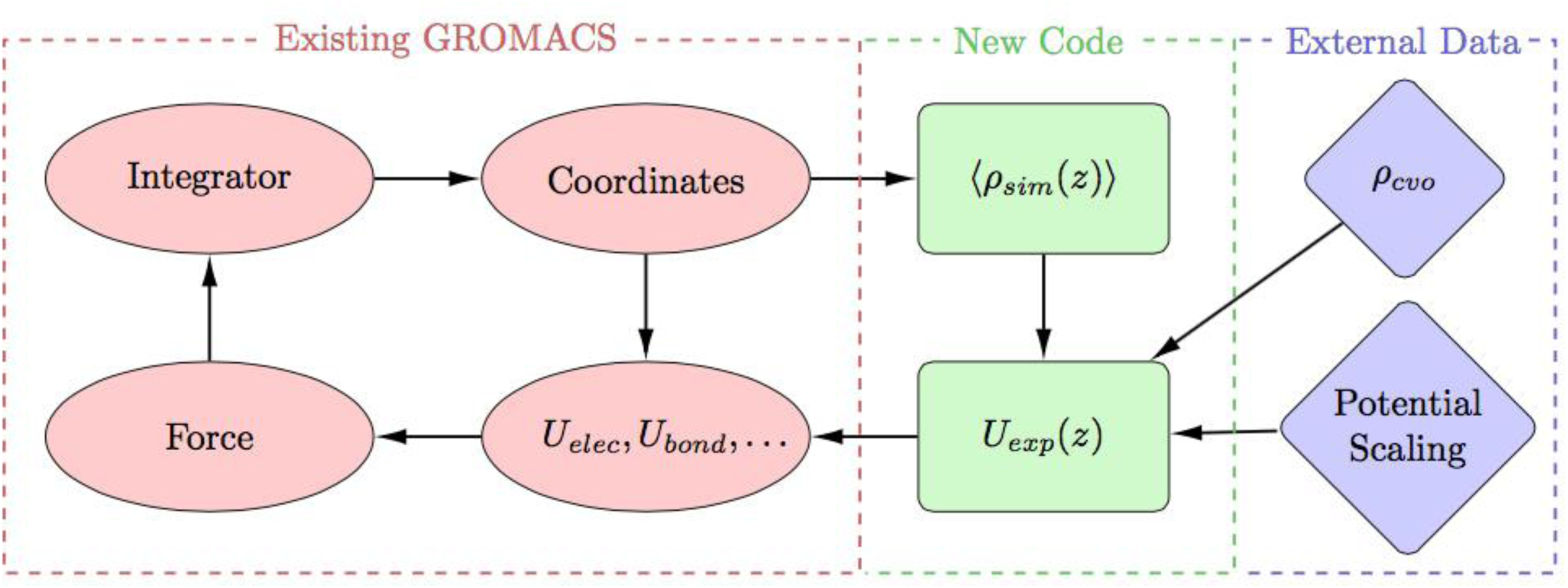
A flowchart for the incorporation of bias into the simulation scheme. The code utilizes atomic coordinates to calculate observables associated with external data, provided before runtime, and constructs a bias which is added to the interactions that determine the system dynamics.

Since 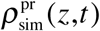 changes as the simulation progresses, *U*_st_(*z*) is adaptive. In the long time limit, as the simulation adopts a conformational ensemble that approaches the experimental one, 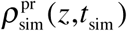 and/or 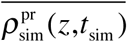 will approach the experimental density. Henceforth, in addition to exploring different settings of *λ*, we use 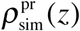 both as a periodically updated instantaneous density or as a time averaged density that accumulates as the simulated trajectory extends.

Polyalanine peptides (*n* = 15 and *n* = 35) were assembled in VMD, placed in a box of (88 Å)^3^ and equilibrated in about 22,400 TIP3 water molecules in 100 mM NaCl under electrical neutrality using GROMACS by first minimizing energy in 50,000 simulation steps, followed by 50,000 simulation steps each in an NVT and an NPT ensemble, and completed with a 10 ns free simulation. An subsequent 10 ns MD run was then performed to create a mock experimental profile *ρ*_exp_, and biased MD simulation runs started with the conformation at the end of this run.

For the PTEN simulations, a POPC/POPS/cholesterol membrane, composed of 800 lipid molecules in total in a ratio of 67:30:3, was built using the Charmm Online Membrane Builder (http://www.charmm-gui.org/?doc=input/membrane.bilayer), and equilibrated as advised.^43^ Protein coordinates were taken from an earlier simulation,^27^ arranged with its center of gravity ca. 30 Å from the equilibrated membrane in a box of 160×160×214 Å^3^ and once again equilibrated in ca. 150,000 TIP3 water molecules with 100 mM NaCl.

## 3. Results

### 3.1 Performance Testing With a Poly-Alanine Model Structure

A 35 residue polyalanine peptide (Ala_35_) that forms stable helical conformations in aqueous solution was simulated to study the efficiency of steering and fine-tune the relative strength of the supplemental potential. A mock experimental profile was constructed from an unbiased MD simulation in solution by averaging densities along the helical peptide axis. The peptide center of mass and moment of inertia from a 10 ns trajectory were aligned to project the density on the helical axis, which in this example defines the *z* axis (Fig. 1A), creating a noisy envelope in analogy to Eq. (2). The resulting profile, smoothed by fluctuations within the helical structure, was used to explore the molecular dynamics of a simulated Ala_35_ molecule under the influence of an external potential that represented an oriented and confined peptide.

We first investigated a 15 residue peptide (Ala_15_) in a series of short MD simulations (typically 10 ns) to select reasonable *λ* values from a wide range between 0.005 k_B_*T*·nm and 2 k_B_*T*·nm. We observed that simulations with bias strengths below 0.05 k_B_*T*·nm were not significantly different from free simulations while bias strengths above 0.5 k_B_*T*·nm suppressed fluctuations strongly. Consequently, we chose a more limited range for *λ* to investigate larger systems in biased simulations. First, we performed biased simulations of Ala_35_ at two different scalings *λ*, set at 0.05 k_B_*T*·nm and 0.5 k_B_*T*·nm, referred to as ‘weak’ and ‘strong’ bias below. We also compared the performances under bias with 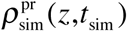 and 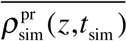 in steering. All simulations were run for 100 ns. In Fig. 3 we show a selection of observables calculated for the biased MD simulation runs. An unbiased simulation is shown for comparison. The center positions do not show a well defined histogram pattern, consistent with the expectation of uniform sampling over all positions at long times. Histograms for simulation runs biased with 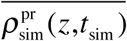 show peaks off-center from the target, with the strong bias giving rise to bimodality. This is because averaged profiles tend to be higher than the target at the center and lower at the fringes throughout the run, drawing the peptide toward the fringes to compensate. The effect is not observed for steering that results from bias with 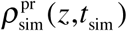. Also, the RMSD histograms match the target well in this case, and the strong bias matches the target particularly closely. RMSDs from simulations biased with 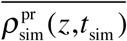 are much broader. Although they start out similar to the instantaneous case, because few samples contributing to the average initially, any mismatch in the profiles that builds up later contributes to the bias and drives the peptide toward regions of mismatch. Since these do not span the entire profile width, the peptide rotates to fill the attractive part of the potential.

**Figure 3:**
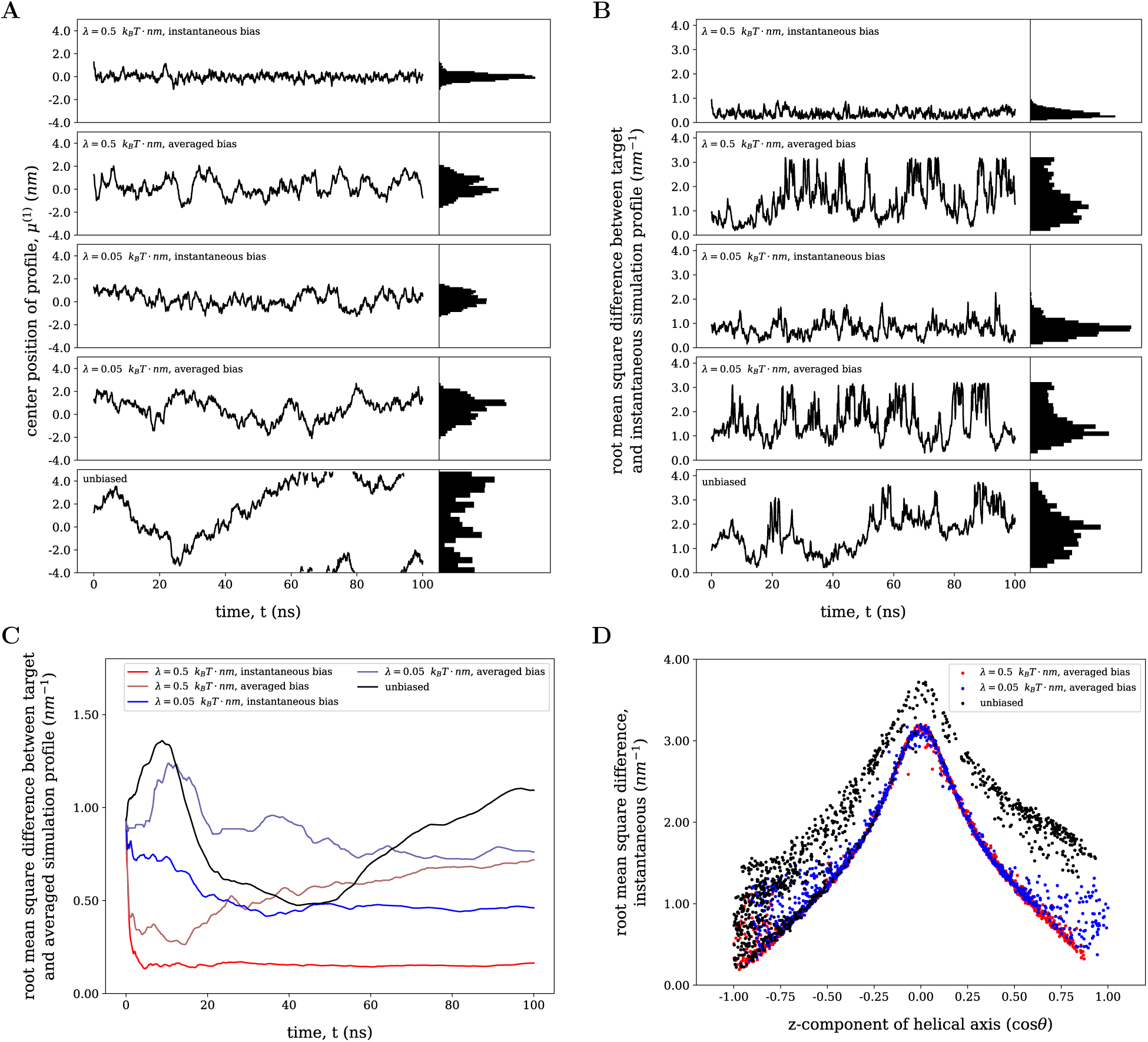
Biased MD simulations of the Ala_35_ helix depicted in Fig. 1A, and their comparison with an unbiased simulation. A target profile is fixed in space with its center located at *z* = 0 Å. (A) Trajectories of the centers of gravity of peptide helices, for different strengths of the biasing potentials and for instantaneous or temporarily averaged profiles, 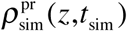 or 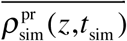, respectively, and their histograms. (B) Trajectories of the root mean square differences (RMSDs) between simulation profiles and target profiles. The truncation of RSMDs near 0.3 Å^−1^ for averages input profiles is due to the algorithm that switches from the biasing potential, Eq. (3), to strong restoring forces, Eq. (6), if fluctuations become s<u>o large that</u> profile overlap is lost. (C) Development of RMSDs between running averages of simulation profiles 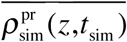 and the target profiles. (D) Instantaneous RMSDs observed in steered and free simulations versus peptide helix axis orientations against the helix orientation of the target which coincides with the *z* axis.

For distinct biasing approaches, Fig. 3C shows how the time averages, 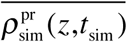, attain steady-state RMSDs from the target density as the simulations progress. While all simulations approach specific long-term values, they do so on quite different time scales. Not surprisingly, the strongly biased simulation biased with 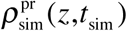 equilibrates most rapidly, within a few nanoseconds, and drops to the lowest steady-state RMSD. This steering approach coerces the simulated object tightly into the target structure. The same steering approach under weak biasing conditions approaches the steady state on a 20 ns time scale and allows for fluctuations which results in a steady state RMSD that is ≈ 3× that for strong biasing. The RMSDs in simulation runs biased with 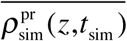 converge more slowly, and fluctuate more strongly, in particular under weak bias. Even if profiles match reasonably well initially, they can lose correlation over time, heading toward long-term RMSD values that are significantly above those that biased with 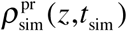. Apparently, the choice of sampling strategy dominates performance, leaving bias strength largely irrelevant, at least bias with 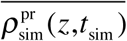.

Figure 3D displays the correlations between the RMSDs of instant profiles and the degree of alignment between the simulated helix direction and the helix direction of the target, which varies between cos*θ* = –1 for antiparallel alignment, zero for orthogonal orientation, and cos*θ* = 1 for parallel alignment. Because the target potential is essentially symmetric, both antiparallel and parallel alignment lead to equally small biasing forces, and fluctuations where the helix axis is nearly orthogonal to the target are strongly suppressed by large biasing forces. Transitions between cos*θ* = –1 to +1 are rare events that occur mostly when the center of gravity of the simulated peptide is near the center of the potential at *z* = 0 Å. Indeed, such transitions are only observed in simulations biased using 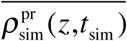 for input which result in relatively large fluctuations, as Fig. 3B shows. Such transitions are virtually never observed in simulations using *ρ*_sim_(*z,t*) as input, where all data points cluster near cos*θ* = 1 (not shown), since fluctuations are suppressed.

To rationalize the overall appearance of RMSD–cos*θ* correlations in Fig. 3D, recall that for off-axis helix orientations (|cos*θ*| ≈ 0), *ρ*_sim_(*z*) is higher and more narrow than *ρ*_exp_(*z*), as shown in Fig. 1, such that it overshoots the target profile where the helix is located and under-fills it elsewhere. Large RMSDs are associated with large local differences between *ρ*_sim_(*z*) and *ρ*_exp_(*z*) which lead to unbalanced restoring forces that result in a torque (Fig. 1D) that drives the helix axis toward alignment with the target (preserving, however, a wrong sign of cos*θ* which is indistinguishable in terms of the density profile). At the same time, the magnitude of RSMDs associated with conformations with |cos*θ*| ≈ 0 do not depend on the position of the helix along *z* as long as the profiles overlap. Once they no longer do, the out-of-range mechanism quickly restores such overlap. As a result, there is little scatter in the RMSDs observed in Fig. 3D for data points near the peak at cos*θ* = 0. This argument, of course, doesn’t apply to unbiased simulations whose RSMDs fluctuate largely, independent of cos*θ*. Conversely, RSMD fluctuations increase near |cos*θ*| ≈ 1 in the conformations (and remain large for the unbiased case) because already small displacements of the profiles’ centers of mass along *z* increase RSMD but restoring force remains moderate or vanishes. Differences in data point densities at cos*θ* ≈ –1 and +1 in the biased simulations indicate that the sampling time (100 ns) was insufficient to allow an adequate number of orientation flips across the region of high biasing potentials near cos*θ* = 0.

### 3.2 Performance With a Well-Studied Protein: The PTEN Lipid Phosphatase

Phosphatase and tensin homolog (PTEN) is a lipid phosphatase whose association with stBLMs *in vitro* has been thoroughly studied with NR^16^ and, independently, with MD simulations.^18,27^ Fortuitously, CVO profiles extracted from experiment and simulation agreed so well that there was little doubt that the MD simulations retraced the experimental results. While the biological significance^44-47^ and structural details^48^ of PTEN are beyond the scope here, the general structure consists of two folded domains (an N-terminal phosphatase domain and a Ca^2+^-independent C2 domain) and a disordered C-terminal tail which accounts for about 13% of the entire molecular weight and controls membrane-accessibility of C2.^49-50^ Here, we test the steering algorithm in MD simulations using NR results obtained for a membrane composed of the phospholipids 1,2-dioleoyl-3-phosphatidylcholine and 1,2-dioleoyl-3-phosphatidylserine with cholesterol as a minor component (DOPC/DOPS/chol = 67:30:3). In the starting configuration, a single copy of the PTEN protein was placed ca. 35 Å away from a pre-equilibrated bilayer^27^ and this system was further equilibrated for 1 ns. Subsequently, all production runs were started under the force according to Eq. (6), because the condition, Eq. (5), applied. They were then either subjected to steering with a potential based on a profile for the protein envelope at the membrane previously determined.^16^ For ‘unbiased’ simulations, the Lagrange parameter *λ*, Eq. (3), was set to zero, but the protein was subjected to the force according to Eqns. (5/6) to pull down to the membrane within the duration of the production run. Based upon our experience with the simulations of Ala_35_ simulations, discussed above, only weak steering potentials (*λ* = 0.05 k_B_*T*·nm) were applied to the protein in biased simulations.

Figure 4 shows trajectories of the RMSDs between PTEN densities that develop in simulations and the one experimentally determined (displayed in Fig. S1C). The PTEN protein approaches the bilayer similarly fast in all three simulations due to bias according to Eqns. (5/6). Subsequently, the RSMDs linger somewhat for the first ca. 15 ns in the unbiased trajectory before the protein attains a stable association with the bilayer. At later times, all three RSMD trajectories settle into regimes that are consistent with bound states of the protein to the membrane, but there remain significant differences between the three simulation approaches. Clearly, both steered simulations show smaller fluctuations of the RMSD than the unbiased simulation, as shown also by the unbiased continuation at *t* > 100 ns in Fig. 4B. However, there are also significant differences in the trajectories biased with 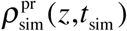 and 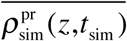 (panels A and B) which require analysis of the underlying profiles and real-space structures.

**Figure 4:**
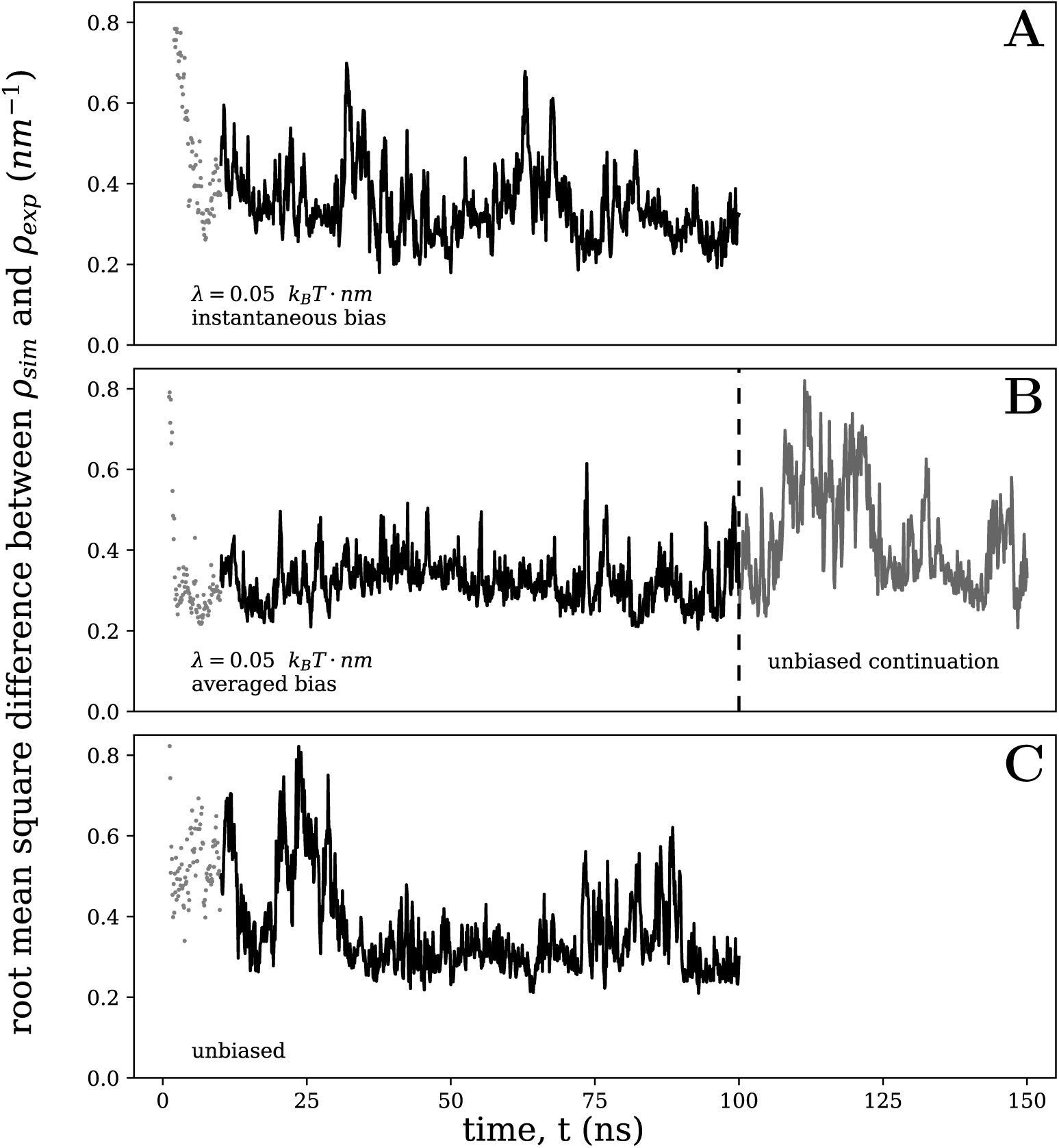
Steered (A, B, *λ* = 0.05 k_B_*T*·nm) and free (C) simulations of PTEN near a DOPC/DOPS/chol bilayer membrane. An NR-determined protein profile of the bilayer (c.f. Fig. S1C) was used to steer simulations by <u>evaluating</u> instantaneous simulation profiles at the time points 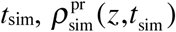, or their accumulating average, 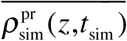. RMS devi<u>ations bet</u>ween simulation and target profiles are shown for simulations bias with 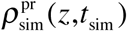 in (A), with 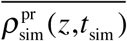 in (B) and without short-range bias in (C), where only long-range attraction according to Eqns. (5 and 6) was applied. Results between *t* = 10 and 100 ns (black trajectories) were further evaluated. The trajectory shown in (B) was continued for another 50 ns after switching off the steering potential, Eq. (3).

Figure 5A shows that all three MD-derived profiles are largely consistent with the experimental result. However, the steered simulation with 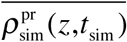 (middle panel) retraces the NR-derived profile better than the other simulations, consistent with the low base level of RMSD in Fig. 4B. Table 1 provides estimates of time points, *t*_*eq*_, in all PTEN simulations where the time series of three observables attain their maxima in uncorrelated samples.^9^ There are apparently large differences in *t*_*eq*_ for different observables. However, only for the biased simulation with 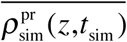 did we observe values well under the runs length for all three observables.

**Table 1:** Convergence times *t*_*eq*_ of selected observables in PTEN simulations with a biasing strength, *λ* = 0.05 k_B_*T*·nm.

| | $\rho_{\text{sim}}^{\text{pr}}(z, t_{\text{sim}})$<br>(‘instantaneous feedback’) | $\overline{\rho_{\text{sim}}^{\text{pr}}(z, t_{\text{sim}})}$<br>(‘averaged feedback’) | free simulation |
| --- | --- | --- | --- |
| mean position ($\mu^{(1)}$) | > 90 ns | 15.5 ns | 29.9 ns |
| RMSD | 4.3 ns | 0.8 ns | > 90 ns |
| profile width ($\mu^{(2)}$) | > 90 ns | 22.4 ns | > 90 ns |

**Figure 5:**
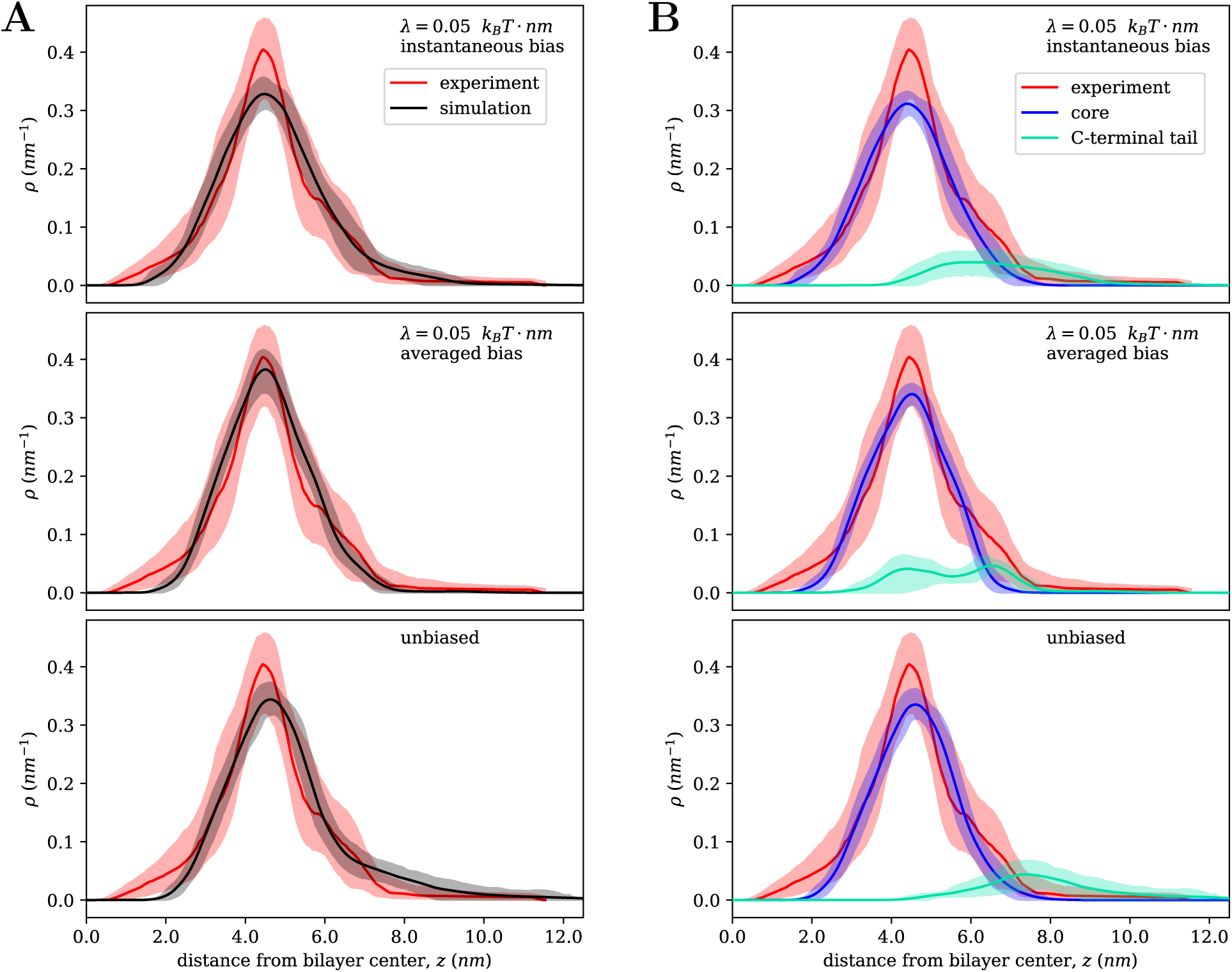
Protein distribution along *z, i.e.* CVO profiles, for PTEN near a bilayer membrane from NR experiment and simulations as in Fig. 4. Median profiles in the normal direction of the bilayer *z* are shown with their 68% (1σ) confidence intervals for the experimental model and the range about the median in the simulations that contains the 68 percentile for density fluctuations (shaded). The origin of *z* is at the bilayer mid-plane. The relation of the protein profiles to the overall structure is shown in Fig. S1. Two biased simulations (*λ* = 0.05 k_B_*T*·nm) and one unbiased simulation were run for 100 ns each, and the trajectories between *t* = 10 and 100 ns were used to determine the profiles. (A) Direct comparison of overall profiles from biased simulations <u>with steering</u> potentials determined by the instantaneous profile 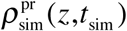 and its accumulating average 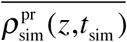 (top and middle panel, respectively), and from an unbiased simulation (bottom panel). In (B), the overall protein density from simulations are decomposed into contributions of the two folded core domains and the disordered C-terminal tail of the protein. This decomposition reveals that the steered simulation with accumulating average profile as the input identified a new tail conformation that leads to a better overall interpretation of the experimental result.

A close inspection of the profiles in Fig. 5A shows that the narrow center of the experimental profile is well captured only in the middle panel. The other two profiles lack density around the central peak, have excess density at high *z* values, and the profile from the unbiased simulation peaks at a slightly higher distance from the bilayer surface than the experimental one. A more detailed analysis, shown in Fig. 5B, reveals the reasons for these subtle differences if we separate the contributions of the folded PTEN domains and its disordered C-terminal tail in the simulation profiles. Because the protein was homogeneously hydrogenated in the NR experiments (no deuteration), these protein regions could not be distinguished in the experimental results. However, in our earlier work, we inferred from an incomplete fit of the crystal structure into the experimental profile that the C-terminal tail is primarily localized at the membrane-distal side of the folded, membrane-bound protein.^16^ This conclusion was subsequently supported by unbiased MD simulations^27^ and the unbiased simulation reported here confirms this finding: C-terminal tail residues that are broadly distributed away from the membrane between *z* ≈ 60–100 Å from the bilayer mid-plane provide conformations that describe the experimental profile reasonably well, even if slight discrepancies between the profiles remain. In physical terms, one may rationalize these protein conformations by observing that the tail carries an excess of acidic amino acids which that lead to its repulsion from the anionic membrane surface.^51^ The simulation biased with 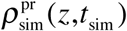 (top panel in Fig. 5B) shows a similar distribution of the C-terminal tail. To enhance overlap with the experimental profile, the the partial profile attributed to the folded domains is slightly broader than that in the middle panel, presumably by enforcing a slightly different tilt angle of the protein body against the membrane.

The steered simulation with 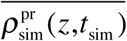 reveals the possibility of a distinct tail organization, as observed from its component profile in the middle panel of Fig. 5B. The associated structural model is the only representation that captures the width of the experimental profile well. Here, the tail profile appears bimodal, a feature that is not observed in the other simulation modes. Figure 6 shows a representative snapshot of the PTEN protein on the membrane in which the C-terminal tail is highlighted. In this model only the initial stretch of PTEN’s tail is dislodged distal from the membrane – corresponding to a peak in the time-averaged profile displayed alongside the structure – while its long C-terminal end wraps around the folded core without interfering with the membrane-binding surface of the protein. Distinct from the simulations in other modes, the folded domains are slightly inclined with their height axis tilted away from the *z* axis, which results in a slightly narrower profile of the protein body along the membrane normal. In combination, these variations of the structure match the experimental result slightly better than the conformations that dominate the other simulation modes and increase the agreement between the profiles to an extent where there is full overlap between the experimental 1σ confidence limit and the breadth of fluctuations of the simulated profiles. Conformations such as that shown in Fig. 6 were not observed in significantly longer conventional all-atom MD simulations of PTEN on lipid bilayers.^18,27^ However, a similar configuration was obtained in the unbiased continuation (grey trajectory in Fig. 4B) between *t*_sim_ ≈ 120 and 150 ns where a comparably small RMSD value between the profiles is observed. This suggests that the alternate tail conformation is not artificially induced by the bias on the one hand and on the other also hints that it may only be marginally stable compared to other tail conformations.

**Figure 6:**
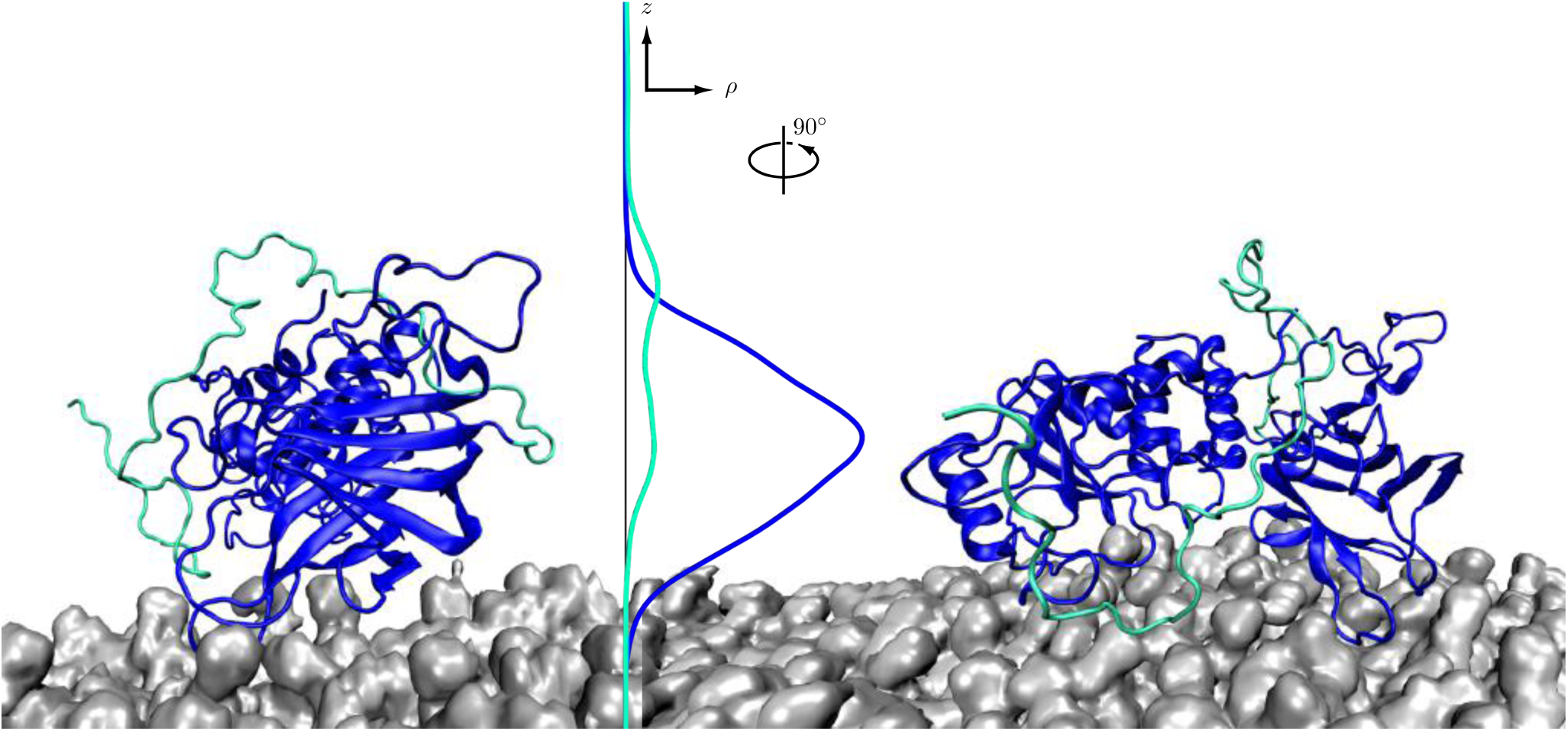
Two orthogonal views of a s<u>imulation s</u>napshot of the PTEN protein on the bilayer membrane at _sim_ = 100 ns (*λ* = 0.05 k_B_*T*·nm, biased with 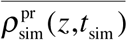). The initial 352 amino acids that contain the folded protein core are shown in blue, the disordered C-terminal tail in green. This color scheme is also applied to the component density along *z*, shown on the right, averaged between *t*_sim_ = 10 and 100 ns.

Figure 7 illustrates the differences between the profiles in the simulation shown in Fig. 4B under bias and after its removal. The top panel shows the overall profile in each segment of the simulation along with the mean profile and the confidence limits from experiment. In the bottom panel the simulation profiles are separated into core and tail contributions. After release of the restraining bias, the protein shifts slightly away from the membrane in the continuation run. Its profile flattens slightly in the center and broadens overall, which reduces its overlap with the experimental result (Fig. 7A). Importantly, the novel, bimodal configuration of the unstructured C-terminal tail obtained in the biased simulation, is retained during the unbiased continuation run (Fig. 7B), suggesting that it occurs in a local minimum of the overall, bias-free potential and thus may correspond to a stable or meta-stable configuration. Because of the low intrinsic resolution of the experiment, such minute overall differences are entirely indistinguishable in interpretations of NR results, and on the basis of MD simulations it cannot be determined which conformation is more relevant for an equilibrium ensemble of structures. Nevertheless, it is important to be able to identify such alternate molecular configurations. In the case of the PTEN phosphatase, the two different tail conformations would lead to distinct conclusions about how the phosphorylation of a serine/threonine cluster (S380, T382, T383, S385) enables cellular control of PTEN binding to the membrane.^49,52^ If nothing else, this example suggests that judicious biasing of membrane-bound protein structures with experimental results from NR increases the discovery potential of MD simulations.

**Figure 7:**
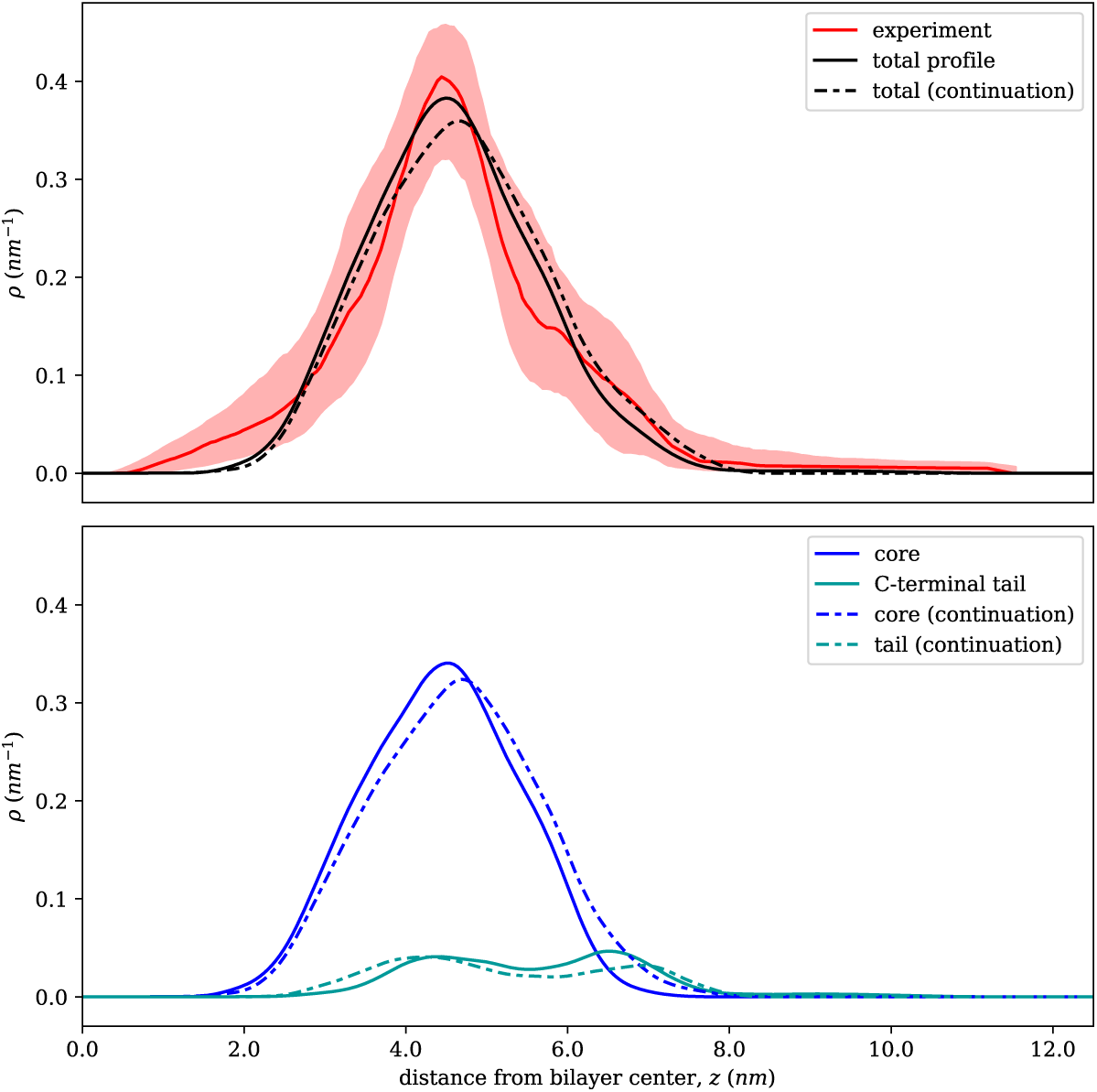
Comparison of the PTEN profiles from the simulation biased with 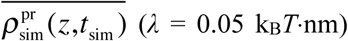, solid lines, averaged between 10 and 100 ns) and from the continuation after removal of bias (dotted lines, averaged over the remaining 50 ns). The top panel shows the overall profiles overlaid on the experimental target with confidence limits. The lower panel separates the overall profiles their into core and tail contributions.

## 4. Discussion

The basic hypothesis underlying our novel approach to steer MD simulations of proteins in or near membranes with results from neutron reflectometry is that the resulting configurations reproduce 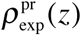 at least as well as those from free simulations while reaching equilibrium properties of the system faster. Examining the RMSD plots in Fig. 3A, B shows that biased simulations of Ala_35_ are indeed more frequently in close agreement with 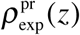 than the unbiased simulation, and the same conclusion is reached for the PTEN results presented in Figs. 4 and 5.

In addition, we investigated whether structural ensembles reach equilibrium faster in steered than in free simulations. Table 1 indicates that steering indeed speeds up equilibration of certain parameters, in particular when determining the biasing potential from 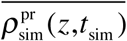. The difference in behavior of the two versions of bias can be rationalized as follows. The bias with 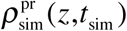 still suppresses fluctuations once an approximate agreement with the experimental data has been achieved, regardless of whether the agreement is true (*i.e.*, the molecular conformation captures the underlying set of structures well) or fortuitous (the molecular conformation is distinct from the true structure but happens to have a similar CVO profile). In distinction, bias with 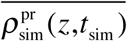 is more lenient in allowing fluctuations, as observed in Fig. 3, which permits the sampling of alternate structural motifs, as inferred from our PTEN simulations presented in Fig. 4 and 5. Fluctuations are not only important to reproduce ensembles representing data from a group of structurally distinct conformations over which an experiment averages. They also explore larger regions of conformational space. As *t*_sim_ progresses, the time-averaged density 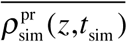 will approximate *ρ*_exp_(*z*) progressively closer as the system samples more conformations that fit the target well. This diminishes the biasing forces and eventually, the system dynamics evolves as if unbiased, and only significant fluctuations away from *ρ*_exp_(*z*) over a substantial time span will affect 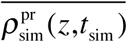 to an extent that a corrective potential emerges again.

Simulation results of the Ala_35_ peptide model, Fig. 3, primarily showed that our approach works well in suppressing fluctuations as expected, depending on potential strength and the dynamics of sampling *ρ*_sim_(*z*). In addition, it demonstrated that for molecules which are not spherically symmetric, the approach can develop torques that align the steered components with the target, despite the fact that the experimental input is strictly one-dimensional. The application of the method to the PTEN phosphatase in a membrane complex provided genuinely new insights. In previous unbiased simulations of this system,^27^ the protein was also swiftly attracted to the membrane surface by the intrinsic interactions. But Table 1 indicates that equilibration is considerably sped up by the steering potential, in particular under bias with 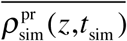. Importantly, this mode of biasing led to the evolution of conformations in the disordered region of the protein that afforded a better fit with the experimental data than in our earlier work. Thereby, using 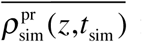 for the determination of the biasing potential is superior in performance than using 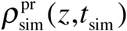 in a number of aspects. While it remains effective in suppressing large deviations from the experimental target profile, it accomplishes this with ‘softer’ steering potentials than instantaneous biasing. This approach thereby allows enough conformational fluctuations that help explore conformation space more efficiently than biasing with 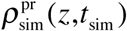, which in turn yields a more direct response to deviations from the target. As in previous work that discussed biasing methods toward other experimental observables,^6-7^ the progressively accumulating form of bias investigated here avoids arbitrary disturbances of the system, and therefore is the most relevant mode of biasing a molecular structure toward neutron reflection results.

## Supporting information

Supplemental file to Manuscript

## Associated Content

Background on the method of neutron reflectometry is provided as a supplement.

## Acknowledgments

This research was supported by the National Science Foundation through XSEDE grants (TG-MCB180120 and TG-MCB190007) and the U.S. Department of Commerce through an MSE grant (70NANB17H299). BWT was supported by a HERE Graduate Student fellowship at Oak Ridge National Laboratory.

## Footnotes

* Note that the values of *A* and *V* do not matter in practical terms as ∫*ρ*(*z*)d*z* is set to unity for both the experimental and simulated densities to determine the shape of the steering potential, which is then scaled with a tunable parameter, *λ*.

