## Supplemental file to Manuscript for "Steering Molecular Dynamics Simulations of Membrane-Associated Proteins with Neutron Reflection Results"

submitted to *J. Chem. Theor. Comput.* version: Sep 3, 2019

#### Background: Reflectometry and MD Simulations

NR is a scattering technique that takes advantage of the stratified nature of a sample, from which an incoming beam is specularly reflected, to provide structural information about the interfacial architecture on the molecular length scale. Here we discuss sparsely-tethered bilayer lipid membranes (stBLMs)[1] in which incorporated phospholipids remain in-plane fluid,[2] providing a suitable matrix for studies of membrane-associated proteins.[3] But this particular sample format is only used as a convenient example: scattering in reflection mode, using both x-rays or neutrons as a probe, can interrogate a large variety of sample formats.[4-11] An stBLM comprises a phospholipid bilayer that is supported by a Si wafer covered with an ultra-flat gold film to which it is tethered via thiol-terminated poly(ethylene-glycol) lipids whose hydrophobic chains intercalate its hydrophobic core. The tether lipid is sparsely distributed on the surface after competitive adsorption with a thiol-labeled spacer (mercaptoethanol, βME). The bilayer is formed around the tether lipids and is separated from the gold surface, passivated by βME, by a hydration layer. Its outer leaflet is free to interact with protein introduced in the adjacent buffer. The resulting supported bilayer structure is complex, but well-characterized.[12]

As with other scattering techniques, specular reflectometry of x-rays or neutrons inherently loses the phase information in the reflected beam, as the measured reflectivity is proportional to the square of the Fourier transform of the interfacial structure.[6] Aided by knowledge of the general membrane structure, well-established parametrization schemes that include auxiliary information, such as chemical connectivity and volumetric information of substructures, laid the foundation for modeling approaches which overcome this limitation.[13-15] Figure S1 illustrates how neutron interference patterns can thus be projected onto the one-dimensional SLD profiles of the interfacial structures (main panel in Fig. S1A and inset). Only the $D_2O$-immersed structures of an as-prepared bilayer and the corresponding protein-loaded bilayer are shown with the tumor suppressor PTEN as an example.[16] Protein binding or incorporation into the bilayer is treated as a perturbation and accounted for by smoothly joining the membrane structure with an atomistic model of the protein from crystallography or NMR. The overall structure is then obtained by adjusting the orientation and penetration depth of the rigid-body structure into the bilayer.[16-19] However, the resulting unique model of the compounded molecular interfacial architecture remains one-dimensional in nature and hinges on the assumption that the protein's crystal or NMR structure persists in its membrane-bound state.[16]

This latter assumption can be critically tested with MD simulations of the corresponding bilayer-protein system. A free simulation of the PTEN protein was set up in a starting configuration where the protein was slightly offset from the membrane. After ≈ 100 ns of the production run, the protein core had settled on the bilayer surface and was further simulated for another ≈ 200 ns.[20] Time-averaged trajectories of the protein on the membrane can now be quantitatively compared with the structure obtained from the reflectometry model (Fig. S1B). For PTEN, the correspondence of the NR-derived and MD-derived structures were striking – and largely fortuitous. Nevertheless, the quantitative coincidence of these structures was interpreted as an independent validation of the neutron model. Moreover, a detailed comparison of MD snapshots of the PTEN core on the membrane and in solution, from an independent simulation, with the crystal structure[21] revealed characteristic differences which were too subtle to be inferred from the NR structure. In addition, the MD simulation suggested that hydrophobic amino acid sidechains associated with the CBR3 loop of PTEN's $Ca^{2+}$-independent C2 domain inserts deeply into the bilayer[20] – a detail that is also experimentally inaccessible, given the resolution of NR. Taken together, such results underscore great potential for linking NR and MD in the atomistic interpretation of protein-membrane complexes, in particular if these techniques are brought into correspondence in a more formalized approach.

### Figures

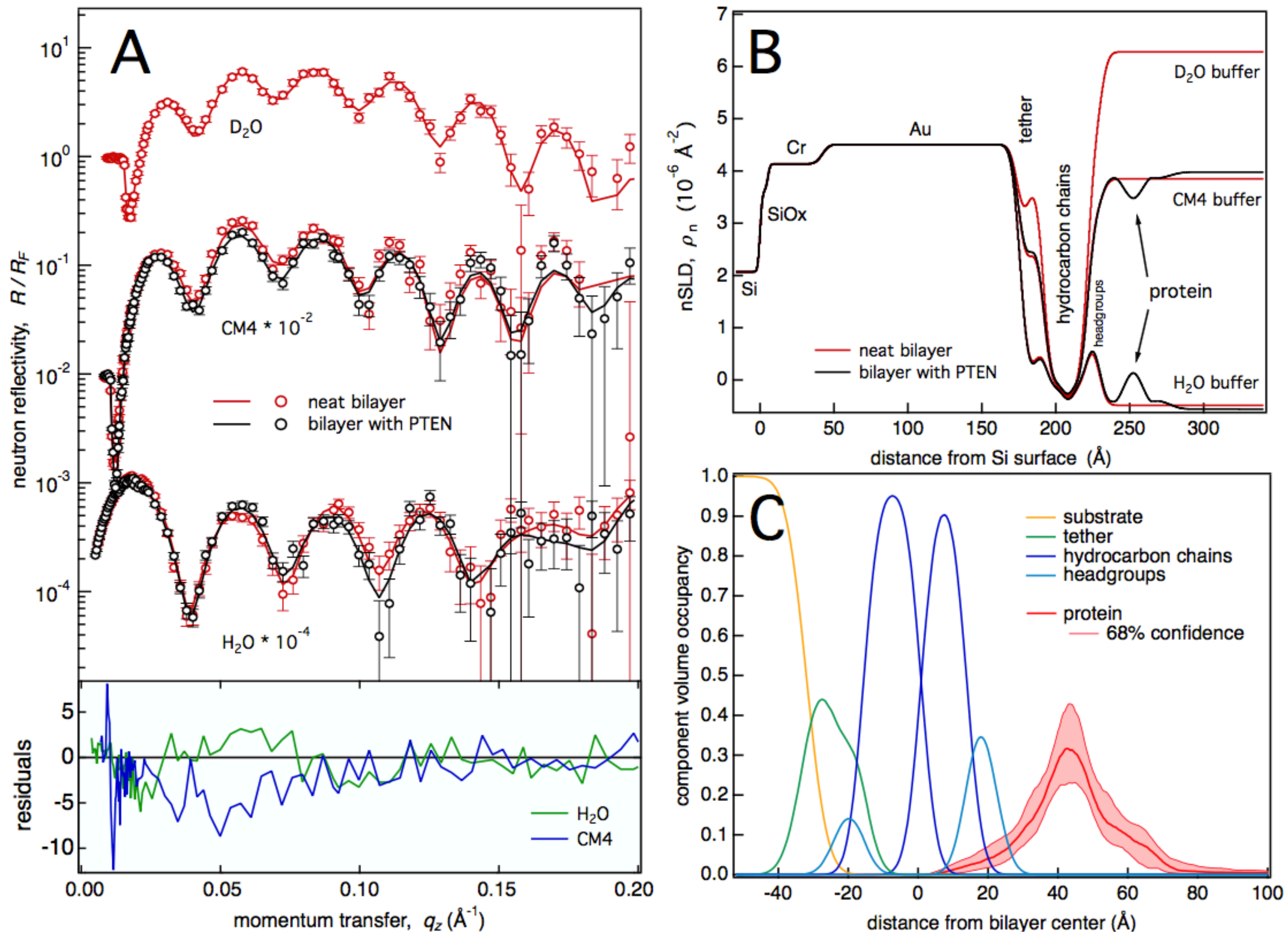


Figure S1: Exemplary neutron reflection data of a bilayer on a gold-coated Si substrate before and after incubation with the PTEN tumor suppressor and their interpretation as a one-dimensional distribution of system components along the interfacial normal direction, *z*. (A) Neutron reflectivities of an stBLM (DOPC/DOPS/chol = 67:30:3) in contact with buffers of different isotopic compositions ($H_2O$, $D_2O$ and a mixture of the two with an SLD, $\rho_n \approx 4\times10^{-6}$ Å$^{-2}$, designated at 'CM4'). Black: As-prepared bilayer. Red: The same bilayer after incubation with PTEN.[16] While the differences in NR with and without protein are small, they are significant, as shown by the residuals plots on the bottom. The modeled reflectivities (solid lines) derive from simultaneous fits to 5 data sets measured under $D_2O$ and $H_2O$ based buffers. (B) Simultaneously optimized SLD profiles derived from a parameterized structural model expressed as a unique component volume occupancy (CVO) profile, shown in (C). The solid lines that fit the data in (A) originate from the SLD profiles shown in (B). The CVO profile, panel C, resolves the complex surface structure that contains the metal films on the Si substrate (at $z < 0$), the tether chemistry and lipid bilayer components of the stBLM (at $0 < z < 40$ Å), and a protein layer (at $z > 40$ Å) in which the PTEN phosphatase skims the membrane surface peripherally. Confidence limits of the protein profile (red) were determined by a Monte Carlo Markov Chain (MCMC) analysis.[22] The algorithms developed in this work use such detailed information to bias MD simulations toward protein conformations that are consistent with the corresponding NR results.
